# Estimating the distribution of reed *Phragmites australis* in Britain demonstrates challenges of remotely sensing rare land cover types at large spatial scales

**DOI:** 10.1101/2023.09.27.559547

**Authors:** Jacob G. Davies, Calvin Dytham, Robert A Robinson, Colin M. Beale

## Abstract

Reed *Phragmites australis* is important for biodiversity, for ecosystem services, and as a resource for humans. Already one of the mostly widely distributed wetland plants globally, reed has recently expanded outside of its native range, modifying ecosystems. However, like most wetland plant communities, reedbed has rarely been mapped at large geographical scales, restricting the information available to ecologists and resource managers. Using Sentinel-2 data and machine learning in open-source software, we produce the first remotely-sensed reedbed map of Britain. A random forest was trained on 79.2 ha of reedbed and 2,719.2 ha of non-reedbed land cover, using free online imagery. Accuracy was high within the training area (AUC > 0.998); however, field validation accuracy was much lower (AUC = 0.671), with many false positives (commission error of 93.4%). A similar workflow carried out in Google Earth Engine, using nearly an order of magnitude more images, gave a lower commission error but a disproportionately higher omission error. Due to the classification error, our map is more useful for a non-spatial estimate of the overall reedbed extent in Britain, rather than for the spatial location of reedbeds in Britain. Using the known commission and omission error, we estimate that in 2015 - 2017 c. 7800 ha of Britain was reedbed. Our study highlights the issues that present enduring barriers to accurate land cover classification at large spatial scales, perhaps suggesting fruitful areas for technological innovation. Even with a ‘big data’ approach and even if technological issues are resolved, ecological factors such as confusion land cover types and geographical variation in temporal reflectance function will probably continue to impose upper limits on the size of area for which land cover can be classified accurately, and therefore on the utility of remote sensing to resource managers.

## Introduction

Global wetland area is rapidly declining, with a net loss of 21% over the last three centuries (Fluet-Chouinard et al. 2023), causing declines in biodiversity and ecosystem services (Clarkson et al. 2013). However, quantitative estimates of wetland area at national scales are scarce (Davidson 2014), with studies often focussing on individual wetlands. To inform policy, for example the initiative to conserve or restore 30% of Earth’s ecosystems by 2030 (Convention on Biological Diversity 2021), there is a need for wetland inventories on a larger geographical scale (Hu et al. 2017).

Reed *Phragmites australis*, a large grass, is one of the most widely distributed wetland plants globally, found on all continents except Antarctica (Packer et al. 2017). Due to its highly competitive ability in specific environmental conditions, it commonly forms large dominant or monodominant patches (hereafter ‘reedbed’). Reed is globally important for biodiversity, ecosystem function, nutrient cycling, and as a resource for humans. Some species (e.g. Eurasian bittern *Botauris stellaris*, southern wainscot *Mythimna straminea*) are almost entirely restricted to reedbeds. Reed expedites ecological succession from open water to land, and plays a complex role as a greenhouse gas source and sink (Brix et al. 2001) and is widely used by humans (Köbbing et al. 2013): for example, for water treatment, biofuel and thatch. In the last century, reed has colonised large areas outside of its native range (Chambers et al. 1999), influencing invasion biology (Meyerson et al. 2016). Due to negative effects, particularly on agriculture, major efforts have been made to eradicate reed outside of its native range (Martin & Blossey 2013). Reed is not globally under threat (Lansdown 2017), but has a dynamic geographical range (Van Der Putten 1997), and so reedbed is often a conservation priority at national scales. To understand its range dynamics and to target its conservation or management, it is important to map reed distribution at large spatial scales and develop tools to allow monitoring of reed distribution over time.

Plant species vary in electromagnetically distinctive characteristics (e.g. concentration of photosynthetic pigments), which themselves vary through the year, and thus can often be distinguished remotely, through classification of spectral reflectances. Additionally, remote sensing especially aids the mapping of hard to access wetland vegetation (such as reed). The training of classification algorithms is aided by reed’s frequent monodominance, easing its identification. Consequently, remote sensing has been an asset to the mapping of reedbed for several decades (Butera 1983). Despite this, few remote sensing studies have mapped reedbed at scales larger than individual wetlands (in which ground-truthing can readily be carried out), with none to our knowledge mapping this wetland type at the national scale. For example, the only estimate of the extent of reedbed in Britain (6,524 ha) stems from a field inventory of British reedbeds made in the 1993 (Bibby & Lunn 1982, Painter et al. 1995). However, those inventories focussed solely on already known reedbeds and so were not exhaustive; no attempt was made to seek out unknown reedbeds. Here, we aim to map the extent of reedbed in Britain completely, using remotely-sensed data. In doing this, we aim to provide a method that can be easily implemented and repeated by ecologists in the future, in order to estimate the change in distribution of this important land cover over time.

## Methods

### Study area, scale and data

We aimed to map all reedbed in Britain, at 10 m x 10 m scale (hereafter ‘10 m scale’). We acquired Level-1C imagery from the Multi-Spectral Instrument (MSI) of the Sentinel-2A satellite (at the time of analysis, the Sentinel-2B/2C sensors had not yet been integrated). Images for all 100 km x 100 km tiles overlapping any of the land surface of Britain and offshore islands except Rockall, captured during the period 1^st^ October 2015 – 30^th^ April 2017, with a given cloud cover of ≤ 5%, were downloaded. The long study period was selected to provide enough non-cloud coverage for the whole study area. GDEM digital elevation model (DEM) data from the ASTER satellite (1 arc-second resolution; ∼30 x 30 m at 55°N) were also downloaded for the study area.

### Pre-processing

Atmospheric correction of satellite data was image-based, and achieved by means of dark-object subtraction, implemented in QGIS (QGIS Development Team 2017) using the Semi-Automatic Classification Plugin (Congedo 2016). More advanced data processing options were not used, in order to implement a workflow that is accessible to those not expert in the field of remote sensing. Reed does not occur subtidally in Britain, and so all tiles were masked to land above the low tide mark.

To identify cloud, a random forest (500 trees, three variables tried at each split) was trained (using all 13 bands) on known cloud/non-cloud for one pass of one scene (30UYD on the military grid) (Liaw & Wiener 2002). Training areas of cloud and non-cloud for this pass were identified visually. This model had an out-of-bag error rate estimate of 1.1%. Cloud presence/absence was then predicted using this model across all scenes, and all predicted cloud pixels were masked out of all images.

Multi-temporal Sentinel-2A images can be mis-registered with respect to each other by up to three pixels at 10 m scale (Skakun et al. 2017), and need co-registering in order to compare by-pixel reflectances over time. Images were co-registered using the *coregisterImages* function in the package *RStoolbox 0*.*1*.*10* (Leutner & Horning 2017). Although the mis-registration was typically eliminated, for some images the function only marginally reduced the mis-registration, or made no improvement at all. Thus it is probably inevitable that our reflectances are slightly spatially smoothed when summarised over time, and our map probably misses some true narrow reedbeds. The co-registration function often failed when the non-NA content of the slave rasters was below 3%, and especially below 1% (e.g. if the tile was mostly sea or cloud). Thus, four scenes for which the non-NA content of the non-master tile never reached above 3% were removed from analysis.

### Multi-temporal images

At any given time of the year, reed has broadly similar reflectance to other vegetation (Gilmore et al. 2008). However, reed’s reflectance and vegetation indices change distinctively through the seasons (Villa et al. 2013), so by using data from more than one season classification uncertainty can be reduced (Onojeghuo & Blackburn 2010).

Ideally, we would have estimated reed’s temporal reflectance function for each band, and used that to predict reed presence/absence; however, data-related and ecological barriers prevented this. Firstly, our cloud classification model could not identify cloud shadow, and so cloud shadows were removed by taking the median reflectance of several cloud-free images within seasons (thereby removing most temporal variation in reflectance from the dataset). Secondly, our study was carried out over 11 degrees of latitude; the temporal function of reed varies over relatively short distances (Haslam 1972, Villa et al. 2013, Tóth 2018). Thus any true reed reflectance function is likely to vary across our study area, creating challenges for its estimation.

We split the dataset into two ‘seasons’; too few cloud-free images were available for some scenes to allow a higher temporal resolution. These seasons were designed to capture periods when reed is generally ‘green’ (‘summer’, June – September) or ‘brown’ (‘winter’, November – April) across the study area. Reed’s ‘green’ season is shorter in the north of the study area (Haslam 1972); and so we left a month between the seasons, to avoid periods when reed might not be universally ‘green’ or ‘brown’ across the study area. The median reflectance was taken for each pixel of each scene for each season respectively.

### Training, classification and prediction

173 training polygons (29 reed, 144 non-reed) with a total area of 2,798.4 ha (79.2 ha reed, 2719.2 ha non-reed) were identified from personal knowledge, Google Maps and Google Street View imagery. These polygons were located on three different scenes (Fig. 1) to train the model on reedbed over a wide geographical area, and on a variety of non-reed land cover types. Edges of training polygons were located away from reed/non-reed boundaries to avoid co-registration errors. Training the classification model using free online imagery avoids one of the two field data collection campaigns associated with remote sensing land cover classification, and the associated financial and time costs.

**Fig.1.**
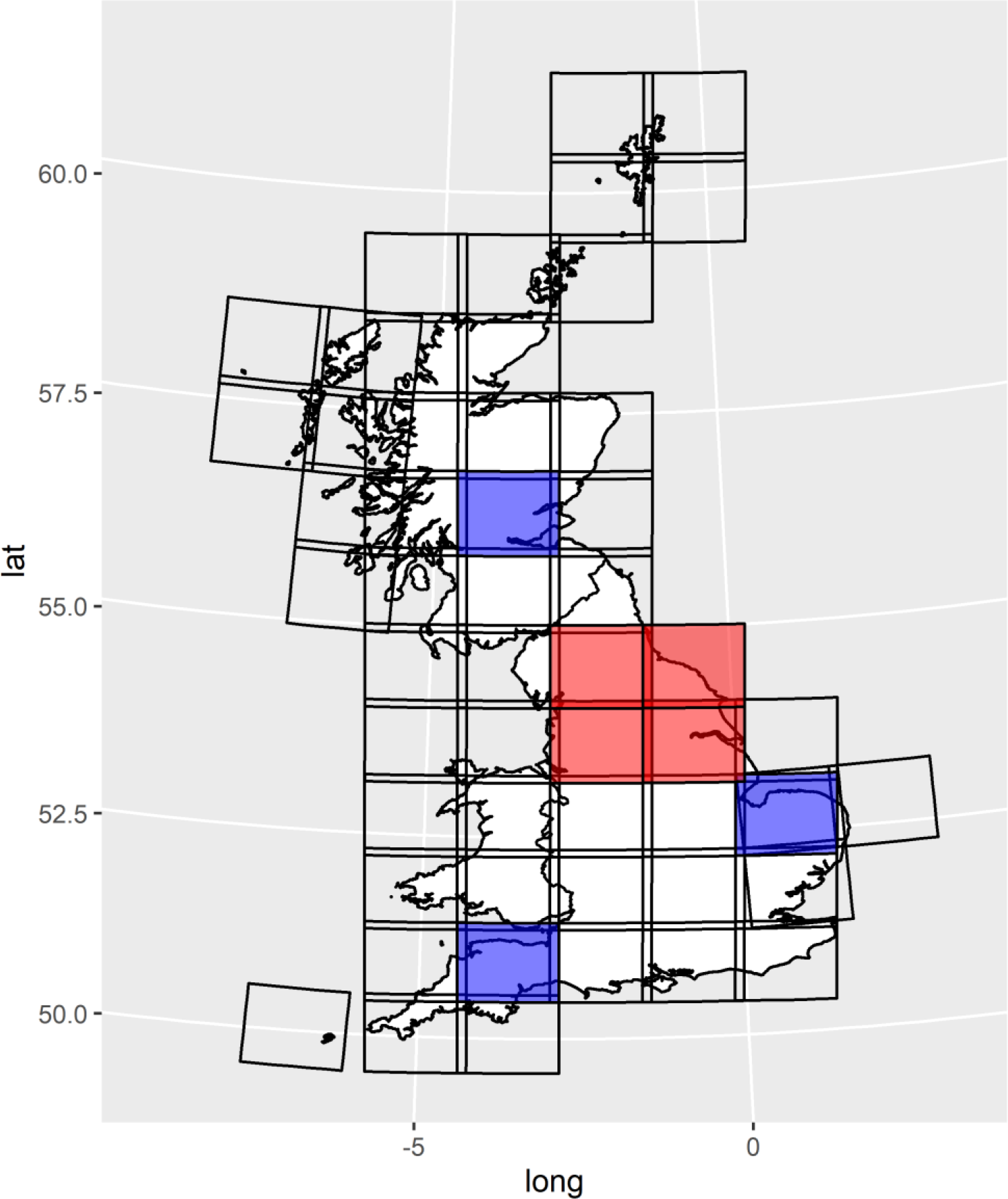
Map of scenes contributing to combined dataset, training scenes (blue) and validation scenes (red).

A random forest (500 trees, three variables tried at each split) was trained on known reed/non-reed. The model was trained at points sampled at random from within the polygons: 100 points from each polygon; points in the same pixel as another point were then discarded. Using band ratios or indices, rather than raw bands, can avoid noise from natural absolute variation in irradiance over multiple dates (Singh 1989), potentially improving classification methods. No single index has proved useful in mapping reedbed across previous studies (Davranche et al. 2010, Villa et al. 2013), so our combined dataset consisted of seven indices, band ratios and standardised bands (Supplementary Table S1) for each season, and the difference in SAVI and NDWI between seasons.

Two studies (Onojeghuo & Blackburn 2010, 2016) found that incorporating texture information improved reedbed classification accuracy. However, those studies used much finer resolution data (2.4 m x 2.4 m pixels) to map large reedbeds. We sought to map reedbeds down to the size of one Sentinel 2-A pixel (10 m x 10 m) – sometimes the full extent of a reedbed – and thus texture measures were unlikely to improve our classification accuracy and were omitted.

Using the trained classification model, reedbed presence probability was predicted across Britain for each scene using the combined dataset. The probability threshold which maximised Cohen’s kappa was calculated for the model: predicted reed presence (*p*) was ‘0’ or ‘1’ if respectively above or below this threshold. Iterative training and re-prediction to improve visual accuracy was carried out until the model could not be improved.

Predicted reedbed maps were aggregated to hectare (100 m x 100 m) scale, before being re-projected to WGS 84 UTM zone 30 and mosaicked together. Ground slope was calculated from the digital elevation model (DEM), and the slope map and DEM were re-projected to the same datum and scale as the predicted reed map. Any cells with a slope of more than 10° or an altitude of more than 470 m (maximum recorded altitude of reed in Britain; Packer et al. 2017) were assigned *p*=0.

### Validation

Covering 11 degrees of latitude (∼1,200 km), it was unfeasible with available resources to validate the map across the entire study area; instead the map was validated across four scenes in northern England (Fig. 1). The validation area was selected to maximise geographical distance from the training polygons and to be central in the study area. Reedbed is a rare land cover type nationally, and thus sampling a random selection of pixels would find too few cells with non-zero probability of reedbed occurrence to estimate either commission error or omission error accurately. Fieldwork was therefore targeted disproportionately towards cells with non-zero probability of reedbed presence. To estimate commission error, 40 hectares were selected from each scene, with 10 randomly selected from within each of the following ranges of predicted proportion (*p*) of reedbed cover from the predicted map: *p* = 0; 0 < *p* ≤ 0.33; 0.33 < *p* ≤ 0.66; 0.66 < *p* ≤ 1. To minimise travel costs, these were selected from the quarter of the scene with the highest non-zero probability of reedbed occurrence.

Each of these hectares was visited and both reedbed (total area of contiguous reed) and reed cover (total area of any reed) of the hectare were visually estimated to the nearest 10%. Then, in each hectare, six randomly selected 10 m cells (three predicted reedbed and three predicted non-reedbed) were visited and reedbed (defined as ≥ 1 m^2^ contiguous reed) and reed presence were recorded. Commission error was quantified in two ways. At the ha scale, predicted and observed cover were regressed, and the coefficient of determination (R^2^) was computed. At the 10 m scale, predicted and observed presences were compared, to give an area under the curve (AUC) of the receiver operating characteristic for each for both reedbed cover and reed presence. Validation fieldwork was carried out from 6^th^ October 2017 to 2^nd^ November 2017.

For the HPC workflow, visual inspection of candidate random forests showed that balanced and unbalanced random forests, and random forests with slightly different training areas, had similar areas of true positives and slightly different areas of false positives. Thus, in order to reduce the area of false positives, two different random forests (‘RF1’ and ‘RF2’) were used for the final model, and the final map was created using the minimum predicted reedbed probability of the two random forests, for each pixel.

All data availability queries, download, classification, raster manipulation and random selection were carried out in R 4.2.1 (R Core Team 2022); pre-processing was carried out in R and QGIS v2.18.

### Workflow in Google Earth Engine

Google Earth Engine (‘GEE’; Gorelick et al. 2016) has become widely used in recent years. GEE uses cloud services to massively scale up computational capability for geospatial analysis, presenting two key advantages over our workflow carried out on a local machine and high-performance computing cluster (hereafter ‘HPC workflow’): much greater data storage capacity and much greater processing speed.

The proportion of Sentinel-2 data that could be incorporated in our HPC workflow was limited by storage capacity. Therefore certain scenes only had a small number of cloud-free passes for a given season, and so the temporal resolution of the data on which the random forest could be trained was limited. We therefore repeated the entire HPC workflow with GEE: to attempt to generate a more accurate reedbed map, and to assess the extent to which relaxing data constraints improves the accuracy of geospatial analysis.

In the GEE workflow we used all Sentinel-2 data (S-2A & B), from the initiation of the Sentinel-2 program (28^th^ June 2015) until the date of analysis (27^th^ July 2019). Temporal resolution was increased from two periods to four: February-March; June-July; August-September; November-December. The combined dataset comprised NDWI, EVI, SAVI, RG, GB, NDVI and SB4 (inter-seasonal differences in SAVI and NDWI were not used, because there were more seasons). GEE imposes user memory limits for tasks, so there was a trade-off between: maximising the number of variables; further increasing the number of images (by increasing the maximum acceptable cloud cover to ≤ 25%); or further increasing the temporal resolution (to eight periods: February, March, June, July, August, September, November and December). One classification model was run for each of these data maximisation approaches, and the accuracy of the resulting map assessed (see validation process below).

The GEE workflow was kept as similar as possible to the HPC workflow. However, some aspects were unavoidably different. Pixels were sampled within polygons (rather than points within polygons), and so it was not possible to balance sampling between categories. It was not straightforward to buffer a raster land mask in GEE, so we used the British shoreline rather than the shoreline plus 100 m buffer: therefore some coastal reedbeds may be missed. The GEE facility to predict probabilities with random forest was not available at the time of analysis, and so presence/absence was predicted. Steep/high altitude terrain was removed from the data stack before training and prediction. We used GEE’s cloud and cirrus removal tools. Because we varied the number of variables between the three data maximisation approaches, we set the random forest to the default setting of number of variables per split: the square root of the number of variables.

The validation fieldwork had already taken place before the GEE workflow was carried out. The validation data were used to assess the accuracy of the GEE reedbed map. The GEE reedbed maps and the validation data are both at 10 m x 10 m resolution, but they have slightly different origins and projections. To assess accuracy at the ha scale, the GEE reedbed map was projected onto the validation map, and predicted reedbed cover was regressed against observed reedbed cover. To assess accuracy at the 10 m scale, the centre points of the 10 m validation cells were re-projected onto the GEE reedbed map, and the predicted and observed presences were compared. As presence/absence (rather than probability) was predicted, AUC could not be calculated for the GEE reedbed map.

## Results

### HPC workflow - structure of random forests

Winter reedbed spectral characteristics (RG, NDWI and GB) constituted two or three of the most informative variables in RF1 and RF2 respectively (Fig. 2). In RF2 the three most informative variables were considerably more informative than the others, but in RF1 there was less separation between these and the remainder. Kappa was maximised at a probability of 0.585 – this was used as the threshold for classifying 10 m pixels.

**Fig. 2.**
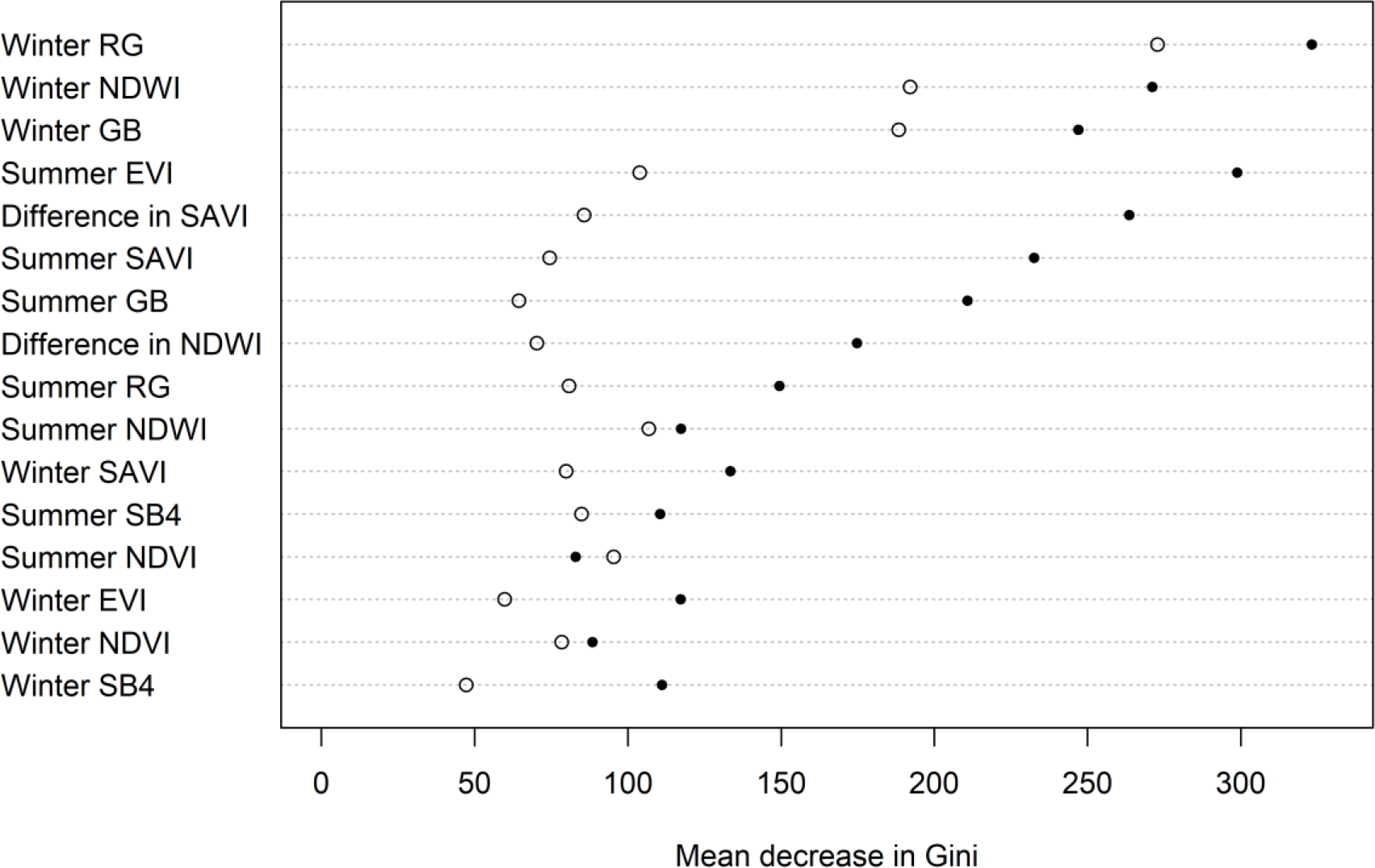
Importance of variables in the two final random forests contributing to the final map (filled circles = RF1; open circles = RF2), HPC workflow. A higher decrease in Gini denotes a greater variable importance.

### Accuracy of HPC reedbed map

Files from 541 passes were acceptable for use after pre-processing (mean 10.82 passes/scene, range 3-22). Each scene had at least one winter and one summer pass. The random forests were predicted over indices derived from these data (Fig. 3).

**Fig. 3.**
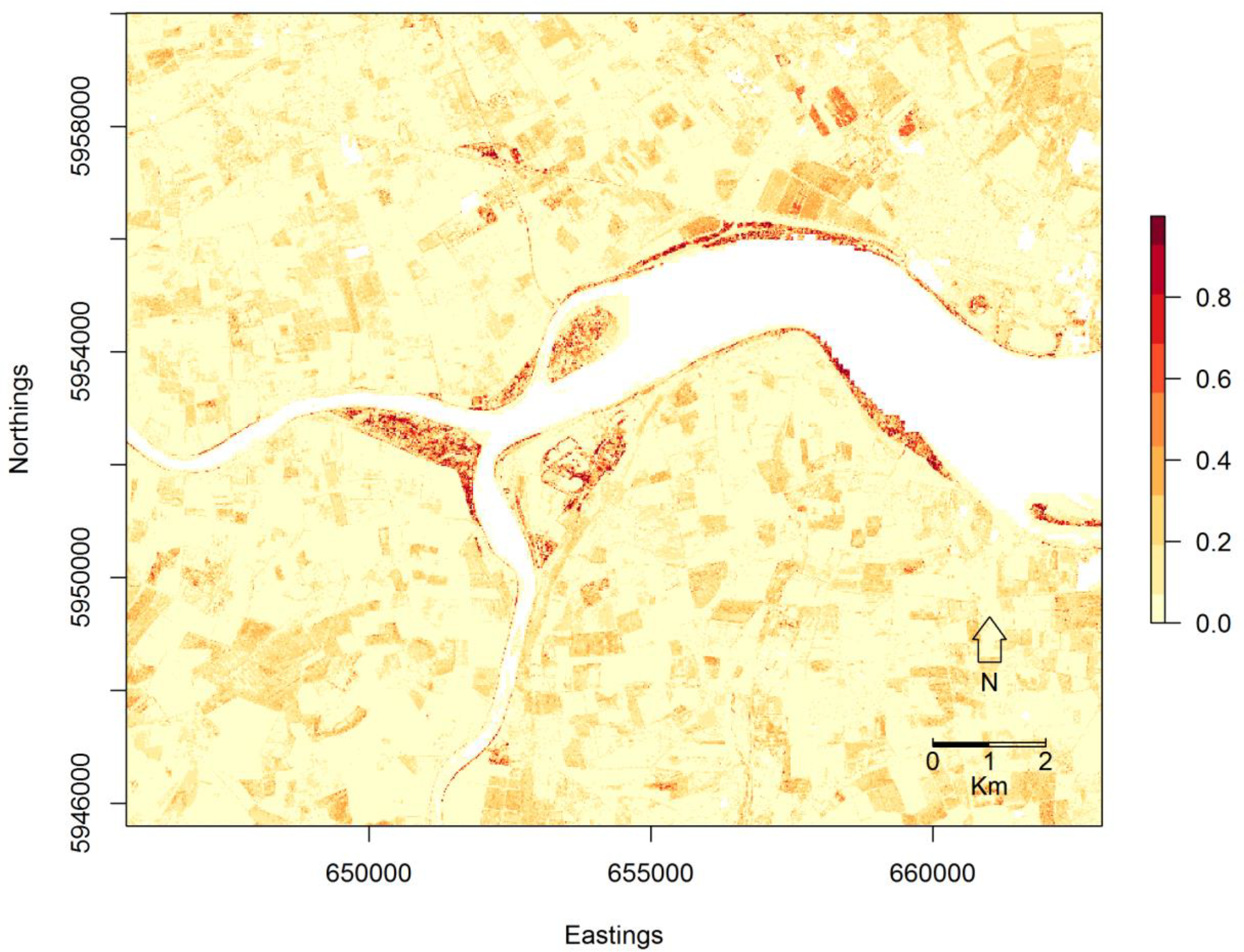
Example predicted reedbed map (colour = predicted probability of reedbed presence at 10 m scale; white = NA) of the upper Humber estuary, NE England, HPC workflow. The dark areas along the edge of the estuary are correctly-classified true large reedbeds.

When tested against the training data, the classification model had near-perfect discrimination: the two random forests used respectively had AUC values of 0.9997 (RF1) and 0.9983 (RF2) against the final map (Fig. 4). Only 154 validation scenes were visited in total, for two reasons: for one validation scene, only eight squares had a predicted probability of 0.66 < *p* ≤ 1; two validation squares from other scenes were not safely accessible. When tested against the validation data, the classification model had much lower discrimination: the combined model had an AUC of 0.671 (Fig. 4, blue line). The overall accuracy of the map at the 10 m scale was 65.1%, but the commission error for reedbed was very high: the majority of predicted reedbed was not true reedbed (Table 1a).

**Table 1.** Confusion matrix for reedbed map at: a) 10 m scale; b) ha scale (proportional reedbed cover converted to binary presence/absence).

| a) 10 m scale |  |  |  |  |  |  |  |
| --- | --- | --- | --- | --- | --- | --- | --- |
| Observed | Predicted |  |  |  |  |  |  |
|  |  | Reed |  | Not reed |  | Omission error (%) |  |
|  |  | HPC | GEE | HPC | GEE | HPC | GEE |
|  | Reed | 21 | 10 | 18 | 29 | 46.1 | 74.4 |
|  | Not reed | 297 | 17 | 588 | 868 | 33.5 | 1.9 |
|  | Commission error (%) | 93.4 | 63.0 | 3.0 | 3.2 |  |  |
| b) Hectare scale |  |  |  |  |  |  |  |
| Observed | Predicted |  |  |  |  |  |  |
|  |  | Reed |  | Not reed |  |  |  |
|  |  | HPC | GEE | HPC | GEE |  |  |
|  | Reed | 11 | 6 | 2 | 7 |  |  |
|  | Not reed | 103 | 10 | 38 | 131 |  |  |
|  | Commission error (%) | 90.3 | 62.5 | 5.0 | 5.1 |  |  |

**Fig. 4.**
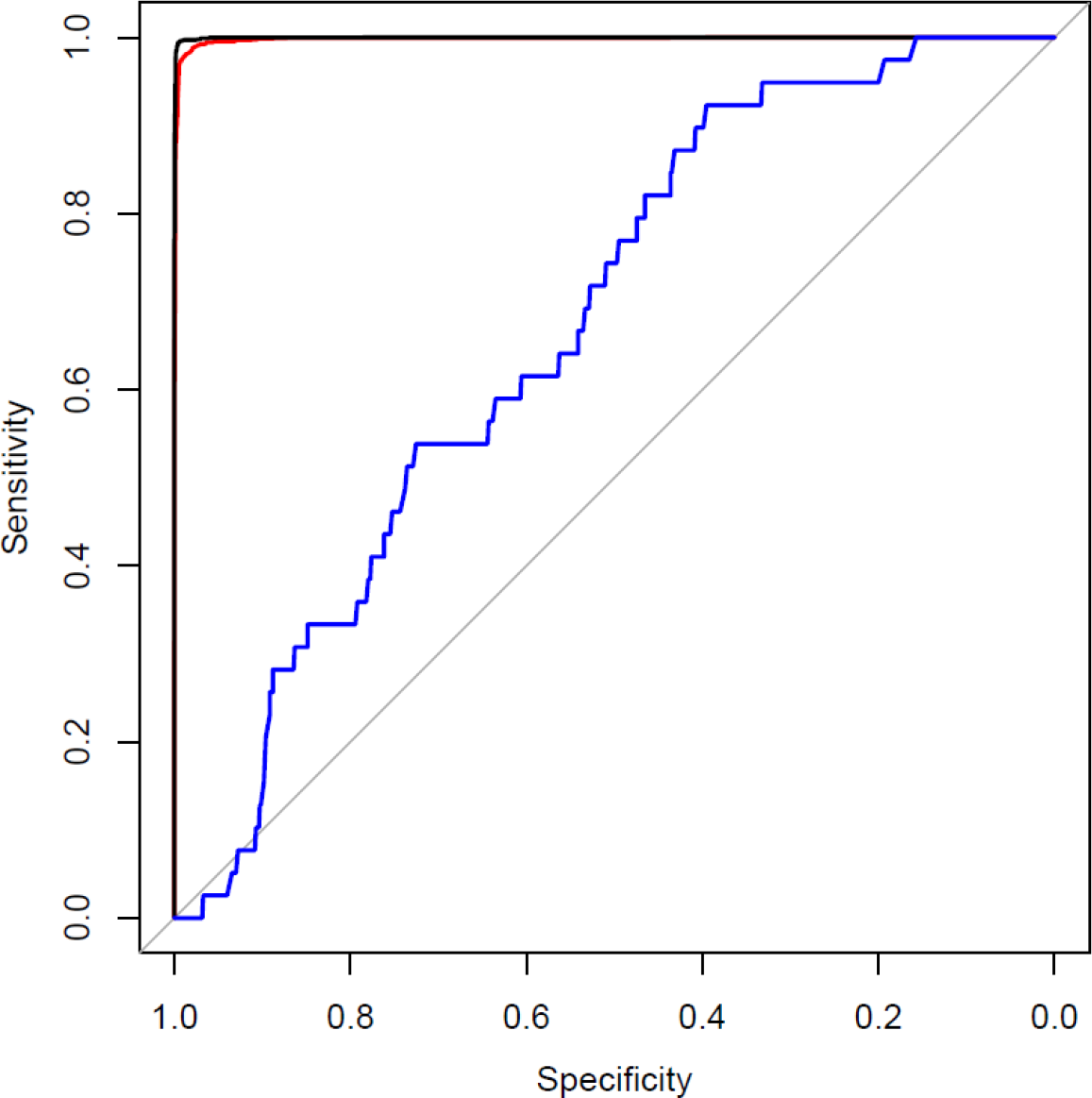
Receiver operating characteristic curve for 10 m pixels, HPC workflow: black, map against RF1 training data; red, map against RF2 training data; blue, map against validation data. Sensitivity = proportion of true positives that are correctly classified; specificity = proportion of true negatives that are correctly classified.

The commission error for reedbed remained very high at the ha scale (Table 1b), although slightly lower than at the 10 m scale. The class frequency of predicted reedbed is deliberately over-represented in our sample (see Methods), and so overall accuracy and omission error are not presented for the ha scale map, because they would be respectively over- and under-estimated. The estimated proportion of reedbed in the landscape was much greater (0.344 at 10 m scale; 0.740 at ha scale) than the true proportion (0.042 at 10 m scale; 0.090 at ha scale).

There was no relationship between predicted and observed reed cover at the ha scale (Fig. 5a). False positives were non-randomly spread among land cover types (Chi-square test, *X*^2^ (1, N = 8) = 32.0, p < 0.0001; Supplementary Table S2). Arable and other open land cover types comprised 61.7% of the sample squares but 73.8% of the false positives.

**Fig. 5.**
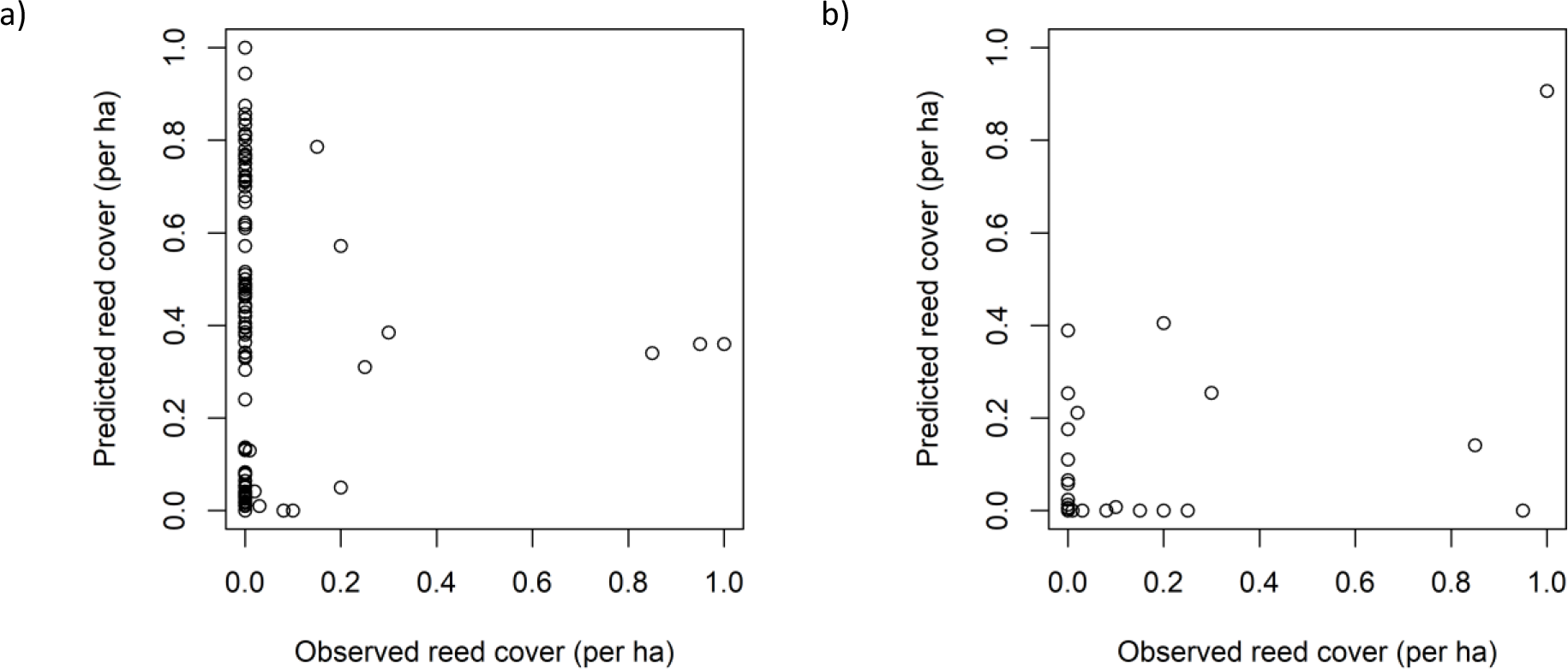
Relationship between predicted and observed ha scale reedbed cover: a) HPC workflow, b) GEE workflow.

### HPC workflow – distribution and extent of reedbed

The distribution of reedbed in Britain estimated using the HPC workflow is presented in Fig. 6. The total area predicted to be covered by reedbed, including known error, is 54,273 ha. Assuming 93.4% commission error and 46.1% omission error, we estimate that 7,765 ha of Britain is covered by reedbed.

**Fig. 6.**
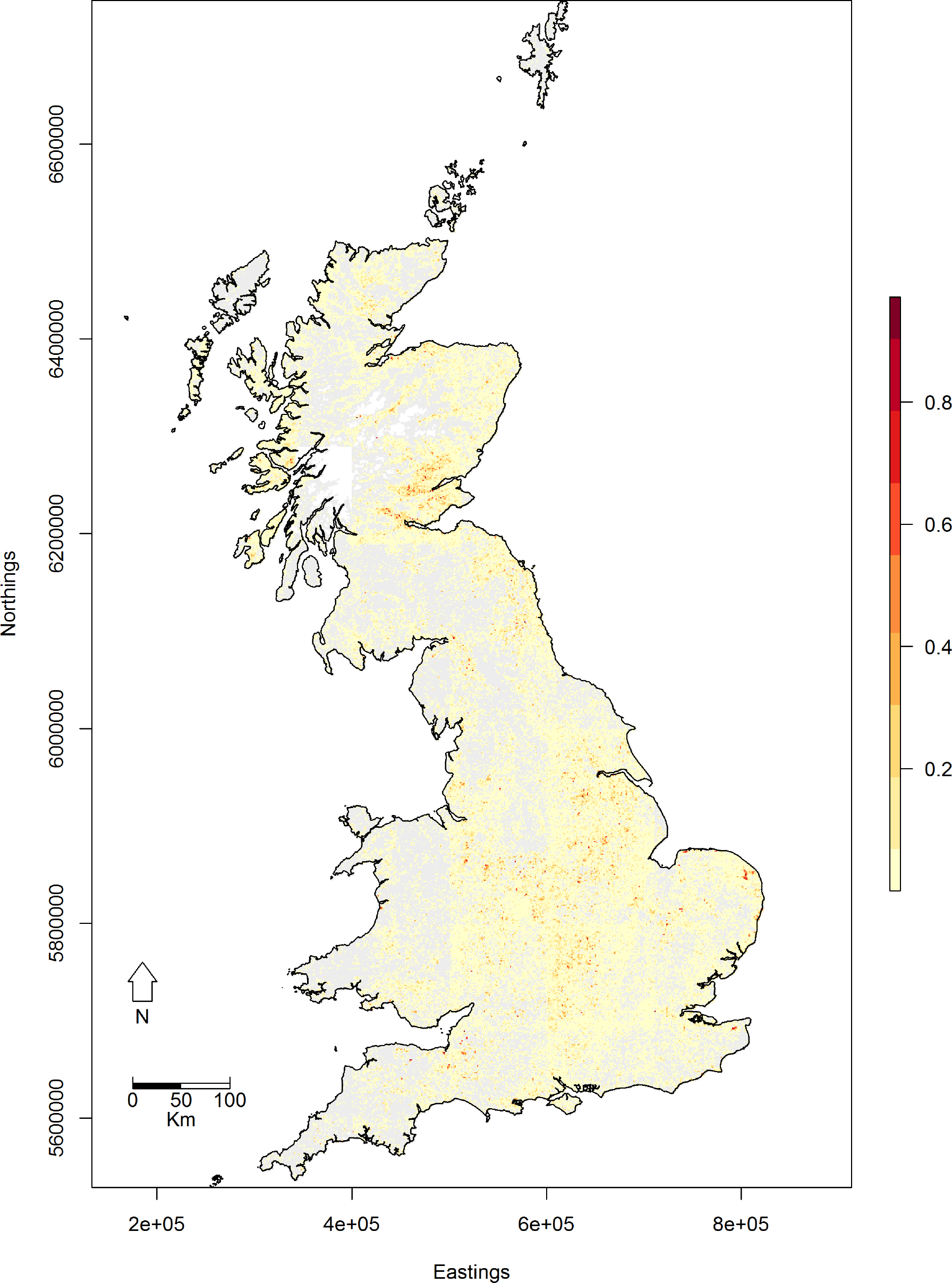
Predicted reedbed map (colour = maximum predicted per-ha cover at 1 km^2^ scale; grey = 0; white = NA) of Britain, HPC workflow. Steep/high altitude terrain has been masked out.

### GEE workflow

It was not possible to balance sampling between categories in the GEE workflow. Therefore only one, unbalanced, random forest was run for each set of criteria. Of the three data-maximisation approaches carried out (maximising number of variables, number of images or temporal resolution), maximising the number of variables gave both the lowest commission error and the lowest omission error: this approach was used. 5,184 images were used, 9.6 times as many as used in the HPC workflow.

At the 10 m scale, the best reed map produced by the GEE workflow had an overall accuracy of 93.8%. This is higher than the overall accuracy of the HPC workflow 10 m scale reed map, but almost identical to the overall accuracy of a map classifying all the validation points as non-reed (94.1%). At the 10 m scale the estimated proportion of reed (0.029) in the landscape was lower than the true proportion (0.042), but at the ha scale the estimated and true proportions of reed were very similar (0.104 and 0.090 respectively). The commission error (63.0%) was still high (Table 1a), but considerably lower than that of the HPC workflow reed map. However, the GEE omission error was higher than that of the HPC workflow reed map: nearly three-quarters of observed reed was predicted not to be reed (Table 1a). The increase in omission error relative to that of the HPC workflow map is proportionately larger than the reduction in commission error.

At the ha scale, the best reed map produced by the GEE workflow had a similar accuracy to the 10 m scale map (Table 1b, Supplementary Fig. S1). There was a weak positive correlation (r = +0.57) between predicted and observed reed cover (Fig. 5b), although this is again likely to be due to the largely correct identification of non-reed, and the high true class frequency of non-reed.

## Discussion

We demonstrate that free remotely-sensed data can be used to predict the presence of a wetland land cover type, with better-than-random accuracy (as assessed by AUC), hundreds of kilometres from the training area. Although field validation was required, the training area itself was only accessed virtually, saving resources. Our reedbed map is the first of Britain and the first to our knowledge at such a large spatial scale. Our estimate (7,765 ha) is the first comprehensive estimate of reedbed extent in Britain, and is of a similar order of magnitude (6,524 ha) to the only other estimate of reedbed extent in Britain (Painter et al. 1995). It is not clear whether the 19.0% increase between the two estimates is due to: a real increase in reedbed in Britain since 1993; the limitation of the previous estimate (Painter et al. 1995) to known reedbeds; or due to error in our classification process.

Remote sensing studies of individual wetlands have typically had high accuracy in identifying reedbed, potentially because of their small geographical scale (Davranche et al. 2010). By contrast, field validation revealed high commission and omission error in our reedbed map. Surprisingly, repeating our workflow in Google Earth Engine with almost an order of magnitude more data from both Sentinel-2 sensors (only data from S-2A were available for the original workflow) and more sophisticated pre-processing methods, did not resolve the accuracy issues. Although the quantity disagreement was reduced in the GEE workflow map, the allocation disagreement was increased. We argue that this lower accuracy in our map stems from the relatively large geographical area covered by our study, resulting in a greater number of confusion land cover types and greater spatial variation in temporal reflectance function.

Our study area, spanning 11 degrees of latitude, covers a wide range of semi-natural and man-made land cover types: much of the UK land area is dedicated to cultivation of grasses *Poaceae* for arable and animal agriculture (19% and 52% respectively in 2019; DEFRA 2020). These constitute a much larger set of land cover types with similar reflectance profiles to reedbed (as evidenced in the high commission error in these land cover types) than is present in a single wetland, making reedbed relatively less distinctive and elevating commission error accordingly. Similarly, reedbed’s relative rarity means that even with a low commission error rate the total area of incorrect commissions would dwarf the total area of correct commissions. This problem is likely to apply generally when classifying other rare and localised land cover types, which are often of disproportionate conservation value.

Reflectance error (e.g. due to instrument error, variation in solar radiation, or cloud removal error) compromises geospatial analysis when the study area is large enough to include multiple scenes. This was evidenced in our study by the visible presence of swath and scene boundaries in the reedbed map (Fig. 6). Massively increasing the number of passes in the dataset (to bring the estimated median reflectance or reflectance-derived measure closer to the true median) with GEE did not resolve this (Supplementary Fig. S1). Reflectance error therefore limits the possibility for predicting outside the swath or scene in which a classification model has been trained.

Reed can vary dramatically in phenology over very small spatial scales (Tóth 2018), and there is spatial variation in reedbed’s temporal reflectance function across our study area (Packer et al. 2017). This potentially makes reedbed’s temporal reflectance function so variable as to overlap with that of other land cover types, preventing a classification algorithm from discriminating between them. This seems to be a key barrier to correctly classifying reedbed at large geographical scales, because increasing the temporal resolution of the data (using GEE) did not improve the accuracy of the map. This issue may also apply generally to vegetation types whose temporal reflectance functions are easily distinguished from those of other vegetation cover over small spatial areas, but which are variable over large geographical areas. This issue could theoretically be resolved by additionally incorporating hyperspectral or non-optical data. However, including SAR data from the Sentinel-1 satellite in preliminary analyses did not improve the GEE map, and no freely available hyperspectral or LiDAR products exist for the whole of Britain. Future analyses could include latitude as a covariate in a temporal reflectance function.

Due to the high classification error, it is more appropriate to use our map (in conjunction with estimated commission and omission error) for a first overall estimate of the extent of reedbed in Britain, rather than for describing the location of individual reedbeds. Repeating our analysis in the future could allow assessment of national change in reedbed extent for natural capital accounting. However, the utility of our map for the latter purpose could be improved by masking confusion land cover types such as arable out with agricultural maps, although this may eliminate some reedbeds which exist as narrow strips in agricultural drains alongside fields.

Classification accuracy of many large geographical scale land cover maps derived from remote sensing is still too low to allow use in a range of applications (Liu et al. 2021). Although repeating our workflow in Google Earth Engine did not resolve the classification accuracy issues, it brought the total analysis time down from several weeks to less than a day, and eliminated the need for large data storage capacity. However, some sources of error remain which are clearly not easily resolved with such a ‘big data’ approach. Classification accuracy for a workflow like ours may improve considerably in the future as satellite reflectance error is reduced, and may improve slightly if other classifiers such as deep learning are used rather than random forests (Venter et al. 2022). However, the number of confusion land cover types and systematic variation in temporal vegetation reflectance functions probably place upper limits on the size of a geographical area that can be classified accurately with such a workflow. Ultimately, this may fundamentally limit the capacity for remote sensing to aid and inform ecological resource management.

## Authorship contribution statement

JD, CD, RR and CB conceived the ideas and designed the methodology; JD carried out fieldwork, analysed the data and led the writing of the manuscript. All authors contributed critically to the drafts and gave final approval for publication.

## Data availability

This study used Sentinel-2 data which are publicly available at the Copernicus Open Access Hub (scihub.copernicus.com) or through Google Earth Engine (earthengine.google.com). Field data used in validation are available at https://github.com/btojacobdavies/GB_reedbed_map.

## Acknowledgements

This research was funded by the Natural Environment Research Council (NERC training grant NE/M009467/1). The funder was not involved in study design, in collection, analysis and interpretation of data, in writing, nor in the decision to submit the article for publication. We are grateful to Rob Critchlow for assistance with Google Earth Engine, and to Gwawr Jones for very useful feedback which improved the manuscript.

