## Supplementary material for "Estimating the distribution of reed *Phragmites australis* in Britain demonstrates challenges of remotely sensing rare land cover types at large spatial scales"

Supporting Information

**Table S1.** Members of the combined dataset, and formulae used to derive them from individual bands (B).

| Quantity | Formula |
|---|---|
| **Enhanced vegetation index (EVI)** | 2.5*((B8-B4)/(B8+(6*B4)-(7.5*B2)+1)) |
| **Green-blue ratio (GB)** | B3/B2 |
| **Normalised difference vegetation index (NDVI)** | (B8-B4)/(B8+B4) |
| **Normalised difference water index (NDWI)** | (B3-B8)/(B3+B8) |
| **Red-green ratio (RG)** | B4/B3 |
| **Soil-adjusted vegetation index (SAVI)** | (1.5*(B8-B4))/(B8+B4+0.5) |
| **Standardised blue band (SBB)** | B4/(B2+B3+B4+B8) |

**Table S2.** Confusion land cover types, HPC workflow. 'Positive' refers to the predicted or true presence of reed.

| Non-reed land cover type | Negatives and true positives | False positives | False proportion of positives |
|---|---|---|---|
| **Arable** | 8 | 51 | 0.864 |
| **Conifer** | 0 | 2 | 1.000 |
| **Deciduous woodland** | 4 | 4 | 0.500 |
| **Freshwater** | 3 | 2 | 0.400 |
| **Grass farmland** | 18 | 14 | 0.438 |
| **Mixed woodland** | 0 | 3 | 1.000 |
| **Not recorded** | 0 | 1 | 1.000 |
| **Other open land cover type** | 11 | 25 | 0.694 |
| **Urban** | 6 | 1 | 0.143 |

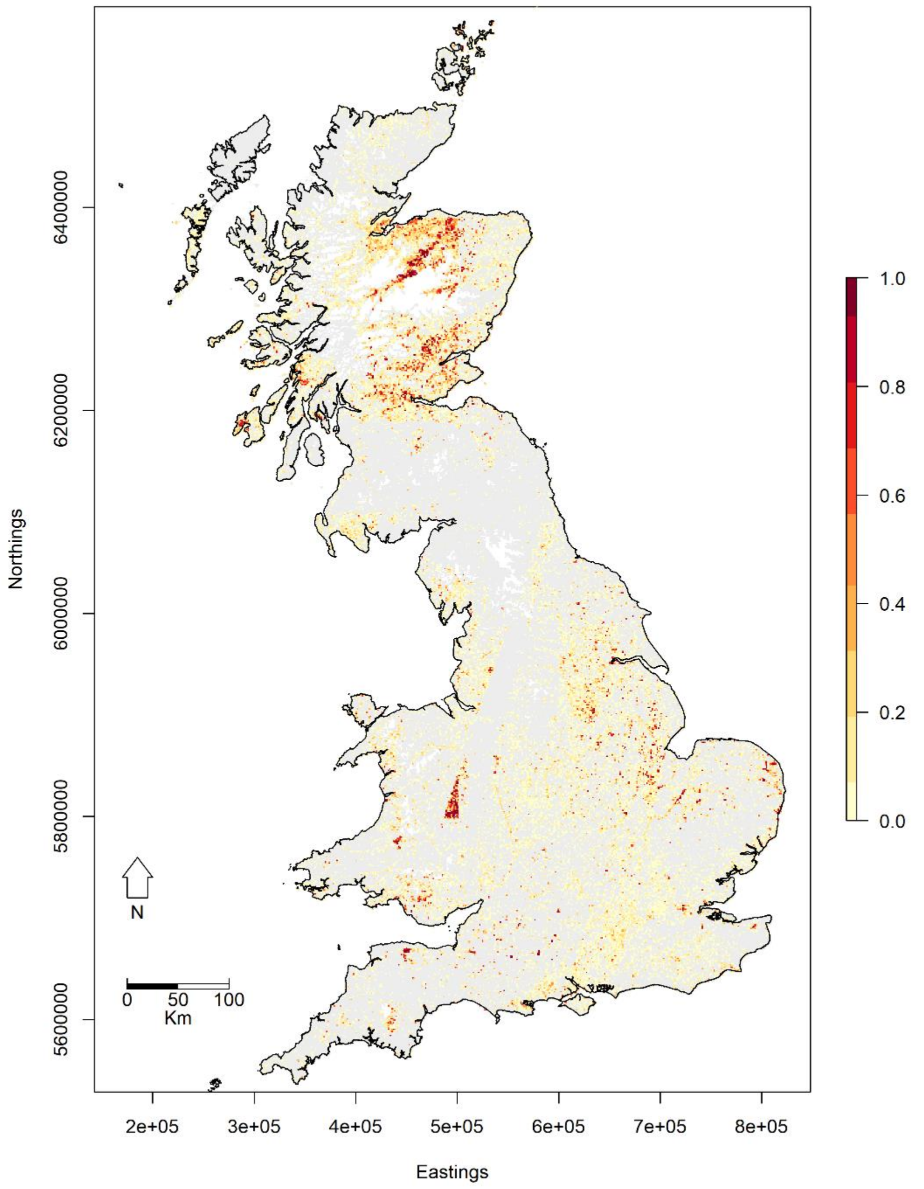


**Fig. S1** Predicted reedbed map (colour = maximum predicted per-hectare cover at 1 km$^2$ scale; grey = 0; white = NA) of Britain, GEE workflow.
